# Phenotypic heterogeneity implements a game theoretic mixed strategy in a clonal microbial population

**DOI:** 10.1101/011049

**Authors:** David Healey, Jeff Gore

## Abstract

Genetically identical cells in microbial populations often exhibit a remarkable degree of phenotypic heterogeneity even in homogenous environments. While such heterogeneity is often thought to be a bet-hedging strategy against unpredictable environments, evolutionary game theory also predicts phenotypic heterogeneity as a stable response to evolutionary “hawk-dove” games, in which rare strategies are favored over common ones. Here we provide experimental evidence for this game theoretic explanation in the context of the well-studied yeast GAL network. In an environment containing the two sugars glucose and galactose, the yeast GAL network displays stochastic bimodal activation. We show that genetic mutants playing the “pure” strategies of GAL-ON or GAL-OFF can each invade the opposite strategy when rare, indicating a hawk-dove game between the two. Consistent with the Nash equilibrium of an evolutionary game, the stable mix of pure strategists does not necessarily maximize the growth of the overall population. We also find that the wild type GAL network can invade populations of both pure strategists while remaining uninvasible by either. Taken together, our results provide experimental evidence that evolutionary hawk-dove games between identical cells can explain the phenotypic heterogeneity found in clonal microbial populations.

## Introduction

Stochastic gene expression is ubiquitous in biological systems ^1–4^. While some noise in gene expression is inevitable, phenotypic heterogeneity is an evolvable trait whose quantitative parameters can be tuned by the architecture and properties of the underlying gene network ^5–8^. This raises the question of what adaptive advantage might be conferred to cells that implement stochastic decision-making ^9,10^. Microbial phenotypic heterogeneity is most often thought to be a response to environmental uncertainty; populations that “hedge their bets” by stochastically adopting a range of phenotypes can gain a fitness advantage if the environment shifts unexpectedly ^6,7,9,11–16^. For example, bacteria may, at some frequency, stochastically adopt a dormant or slow-growing “persister” state, which has reduced fitness when times are good, but which is more likely to survive in the event of catastrophic environmental stress ^12,17^.

Other evolutionary drivers of heterogeneity, such as altruistic divisions of labor and evolutionary “hawk-dove” (or snowdrift) games, are distinct from bet-hedging in that they result from interactions between individuals within populations, and can manifest even in deterministic environments ^18–21^. Altruistic divisions of labor occur when one phenotype sacrifices its fitness to increase the fitness of the remaining population. Canonical examples include the self-sacrificing virulent phenotype of *S. typhimurium* ^2,22^ and colicin production, in which toxin is only released upon cell lysis^23^. Because there exists the potential for an individual to gain a fitness advantage by never adopting the low-fitness phenotype, such altruistic divisions of labor must generally be maintained by inclusive fitness effects such as kin or group selection^15,24^. In contrast, phenotypes in hawk-dove games are mutually invasible. Such games tend toward an equilibrium wherein all phenotypes have equal fitness: either a stable coexistence of pure strategies or a single evolutionarily stable mixed strategy (mixed ESS). Identifying which of these three evolutionary drivers (hawk-dove games, uncertain environment, altruistic division of labor) is at work in a given phenotypically heterogeneous population is complicated by the possibility that multiple of these phenomena can coexist in a given system ^20,25^. While bet-hedging and altruistic division of labor have been observed in microbial populations ^11–13,22,25^, the relevance of hawk-dove games to microbial phenotypic heterogeneity remains largely unexplored.

Evolutionary game theory concerns itself with situations in which the fitness of a phenotype is a function not only of the individual’s own phenotype, but of the phenotypes adopted by other individuals. In the hawk-dove game of animal conflict (Box 1), neither pure strategy (ie “play hawk” or “play dove”) is evolutionarily stable, since populations of each can be invaded by a minority population of the other. Because of this mutual invasibility, if allowed to evolve, the system reaches a stable equilibrium that contains a mix of phenotypes. In microbial populations, such negative frequency dependent interactions have been shown to stabilize the coexistence of different genes ^26–28^.. However, a genetically identical population can theoretically achieve the same stable mix of phenotypes, provided each individual randomizes between strategies with the appropriate probabilities. Such stochastic choices between strategies are called *mixed strategies,* and the specific probabilistic strategy that implements a stable equilibrium is the *evolutionarily stable mixed strategy* (or mixed ESS).

### Box1

#### Box 1: The Hawk-Dove game and evolutionarily stable mixed strategies

Evolutionary Game Theory is the branch of theoretical biology concerned with biological “games,” or situations in which an individual’s evolutionary fitness is a function of the strategies adopted by other individuals in the population.

In evolutionary game theory, an individual’s *strategy* is a specification of what phenotype an individual will adopt in any situation in which it may find itself. Strategies can be deterministic (pure strategies) or probabilistic distributions over the pure strategies (mixed strategies). The central concept in evolutionary game theory is the evolutionarily stable strategy (ESS). An ESS is a strategy such that “if all members of the population adopt it, then no mutant strategy could invade the population under the influence of natural selection.” ^22^ Evolutionarily stable strategies are subsets of Nash equilibria from classical game theory.

Evolutionarily stable mixed strategies (mixed ESS) are a class of ESS that arise in “anti-coordination” games, or games in which rare strategies are favored over common ones. The canonical example of these is the two-player symmetric hawk-dove game as described by Maynard Smith^22^. In this game, two animals are competing over a resource of value *V*, and each animal has two pure strategies available: to fight (Hawk) or retreat (Dove) from a fight. There are three possible pairings in a symmetric two-player Hawk-Dove game:

1. If Hawk meets Dove *(H,D),* Dove flees and Hawk gets the resource (*V*). Dove retreats and receives a payoff of zero.
2. If Hawk meets Hawk *(H,H)* they fight. Each has a ½ probability of getting the resource, but only after incurring an injury cost (*C*) greater than the value of the resource. Their expected payoff is ½ (*V-C*), which is less than the payoff from retreating.
3. If Dove meets Dove (*D,D*) they share the resource. Their expected payoff is ½(*V*)

In matrix notation, the payoffs are as follows (payoffs are to the left player):

|  | H | D |
| --- | --- | --- |
| H | $\frac{1}{2} (V-C)$ | $V$ |
| D | $0$ | $\frac{1}{2} V$ |

Since the expected payoff of playing Hawk against Hawk, *E(H,H)* is less than zero, the payoff of playing Dove against Hawk, it is apparent that the Hawk strategy cannot be evolutionarily stable, since a single Dove will receive a higher payoff and invade the Hawks. Likewise Dove is also not evolutionarily stable, since *E(H,D)* > *E(D,D)*; a single Hawk will invade a population of all Doves. Thus, the pure strategies are mutually invasible. In a population consisting of both Hawks and Doves, if Hawks are present with frequency *f* and Doves with frequency *(1-f)* then for strategy *I*, the expected payoff of adopting *I, E(I)*, is as follows:

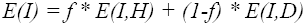

Suppose now that individuals can implement a mixed strategy. We will define strategy *I* as “play Hawk with probability *p*, and Dove with probability *(1-p)*.” The evolutionarily stable mixed strategy (mixed ESS) will be to choose *p* such that *E(H, I) = E(D, I)*. In other words, any single invader will be indifferent to playing either Hawk or Dove (or, consequently, any possible probabilistic combination of the two). Therefore, an isogenic population playing the mixed ESS would be uninvasible by a single newcomer implementing any other strategy, pure or mixed.

It is important to note that while the ESS is uninvasible, it is not the strategy that maximizes the payoff. If *V=1* and *C=3*, for example, the mixed ESS is to play Hawk with frequency 1/3. The evolutionarily stable population receives a mean payoff of 1/3, whereas a population of pure Doves receives the maximum mean payoff of 1/2. Below are plots detailing the payoffs by strategy (upper) and for the overall population (lower), as functions of the frequency of Hawks when *C = 3V*.

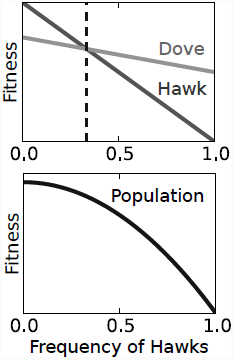

**Table 1.**
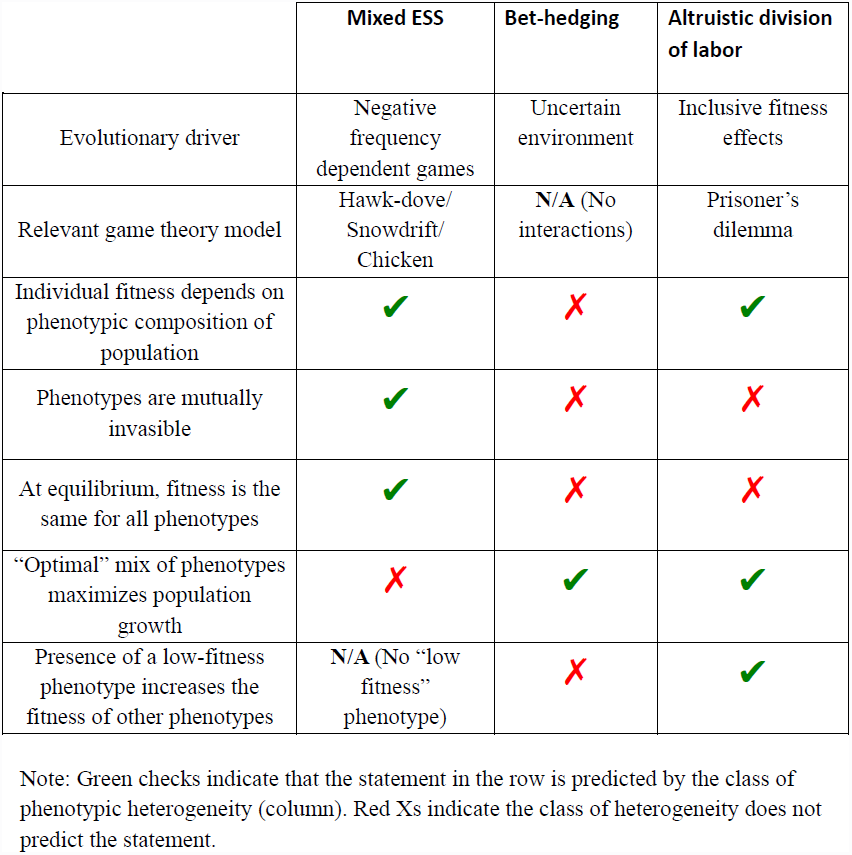
Evolutionary classes of phenotypic heterogeneity

Mixed ESS in hawk-dove games have several experimentally observable characteristics that distinguish them from bet-hedging strategies and altruistic divisions of labor (Table 1). The primary defining characteristic of the hawk-dove game is that the pure strategies, or phenotypes, are mutually invasible: each pure strategy can invade the other when rare. Secondly, if the mixed strategy is evolutionarily stable, all individuals in the population—and potential invaders implementing any other strategy, pure or mixed—will receive an equal payoff ^29^. Thirdly, while other strategies of phenotypic variation maximize some measure of population growth (see discussion and Table 1), evolutionarily stable mixed strategies are not necessarily growth optimal for a population (Box 1)^30^.

### Results

To study stable mixed strategies in the laboratory, we investigated the decision of the budding yeast *S. cerevisiae* regarding which carbon source to consume. Yeast prefers the sugar glucose, but when glucose is limited yeast can consume other carbon sources^31^. The well-studied yeast GAL network contains the suite of genes required to metabolize the sugar galactose. The GAL network activates as a phenotypic switch: each cell is either GAL-ON or GAL-OFF^32,33^. The network is under catabolite repression by glucose^31^; however, yeast can still activate the GAL genes in the presence of modest glucose concentrations provided there is galactose in the media as well. Furthermore, in a wide range of glucose and galactose environments, the GAL network is neither uniformly activated or deactivated across the population, but is expressed bimodally ^33,34^ (Figure S1). In mixed sugar conditions, a tradeoff exists between activation and repression of the GAL network: activation of the GAL network in the presence of glucose may provide some benefits to the cell in consuming galactose ^34^, but expression of the GAL genes also imposes a significant metabolic cost (Figure S2). Similar tradeoffs in catabolite-repressed networks have been characterized previously ^7,35–37^, but have been studied in the context of sympatric speciation or bet-hedging.

GAL bimodality in mixed glucose and galactose suggests a similarity to the following hawk-dove-like foraging game. In this game, an isogenic population is confronted with a phenotypic decision to “specialize” in consuming one or the other of two limited food sources, A and B (Figure 1A-B). The more individuals who adopt the pure strategy “specialize in A,” the more quickly A will be consumed, reducing the payout to individuals who chose that strategy. Hence, if all individuals choose “specialize in A,” an incentive may exist for an individual to choose instead “specialize in B,” and vice versa. The resulting equilibrium consists of a stable mix of the two pure strategies; therefore, an isogenic population that can adopt that stable mix via phenotypic heterogeneity would be uninvasible.

**Figure 1.**
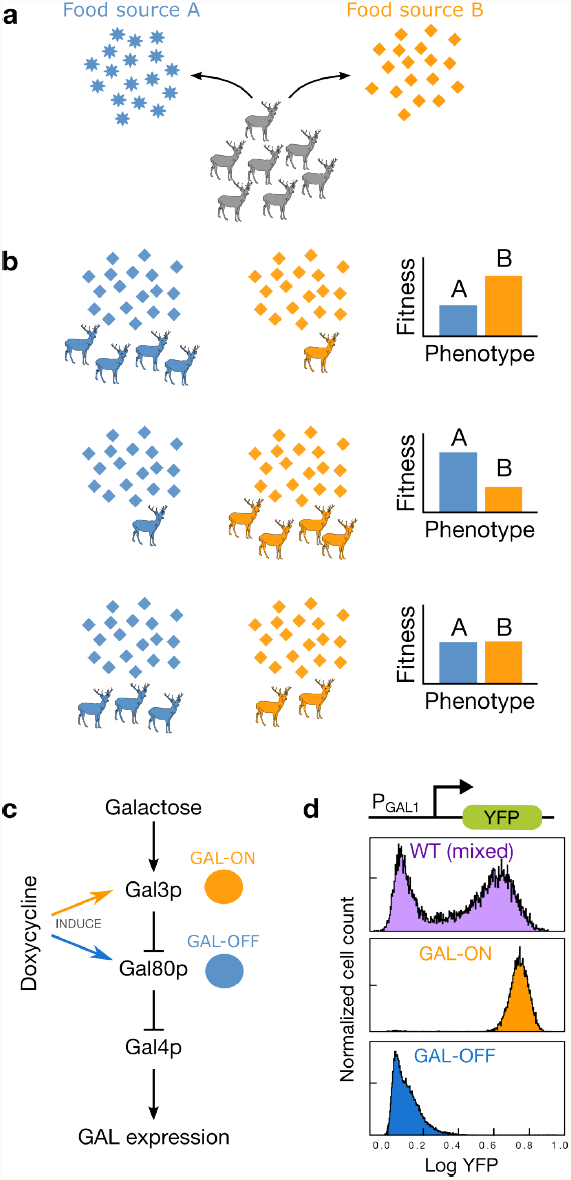
A simple foraging game with multiple food sources can favor phenotypic heterogeneity. **a,** A simple foraging game with a mixed Nash equilibrium. Each member of a group of foragers is confronted with a binary decision about whether to specialize in consuming food source A or B. We assume that individuals choose simultaneously and without knowledge of the actions of others. Resource limitation makes it a game; each individual’s payoff is a function of the actions of other individuals. **b,** If all other members of the population adopt some pure strategy (e.g. “specialize in food A”), an individual opting for the opposite pure strategy (e.g. “specialize in food B”) gains a fitness advantage (top and middle panels). The Nash equilibrium of the simple foraging game is reached when the population divides between the two sources such that both phenotypes receive the same fitness and there is no fitness incentive for any single individual to change strategies (lower panel). In such a game, if each individual adopts the mixed strategy that stochastically chooses between pure strategies with the equilibrium probabilities, then that mixed strategy is evolutionarily stable. Though this game is not necessarily representative of real life foraging scenarios, it serves to illustrate why we might expect environments with multiple food sources to favor the evolution of mixed strategies. **c,** Gene expression in the yeast GAL network is regulated in part by *GAL4, GAL80,* and *GAL3* (full network not shown). A GAL-OFF pure strategist is engineered by inducing the expression of *GAL80*, whose protein product inhibits GAL expression. Likewise a GAL-ON pure strategist can be engineered by inducing expression of *GAL3*, which inhibits GAL80 in the presence of galactose. **d,** In a mixed sugar environment (0.03% glucose, 0.05% galactose), “GAL-ON” and “GAL-OFF” pure strategists remain unimodally activated and inactivated, respectively, while the wild type GAL network exhibits bimodal gene expression. Cultures in Figure 1d were initially grown overnight in 0.01% (w/v) glucose and 1ug/mL doxycycline to saturation, then diluted to an OD of 0.002 and grown 8 hours in mixed glucose and galactose before measuring GAL activation via flow cytometry.

Given the bimodal expression of the yeast GAL network in some conditions, we sought to probe experimentally whether this phenotypic heterogeneity might be the implementation of an evolutionarily stable mixed strategy in response to a foraging game. Since mutual invasibility of phenotypes is the defining characteristic of a hawk-dove game, we began by competing mutant “pure strategists” at many initial population frequencies. As a GAL-OFF pure strategist, we used a yeast strain whose native *GAL80* (a repressor of the GAL network, Figure 1C) was replaced with a mutant version containing a tet-inducible promoter ^33^. As a GAL-ON pure strategist, we used a mutant whose *GAL3* (a repressor of *GAL80*, Figure 1C) was similarly tet-inducible. We confirmed that, in the range of glucose and galactose concentrations that induce bimodality in the wild type yeast, our doxycycline-induced GAL-OFF and GAL-ON pure strategists are unimodally inactivated and activated, respectively, for GAL gene expression (Figure 1D, Figure S1).

To test for negative frequency dependence between the pure strategists, we mixed six biological replicate pairs of RFP-labeled GAL-ON and CFP-labeled GAL-OFF strains at a total of 60 different initial frequencies, and incubated them for 20 hours in a mixed glucose and galactose environment. To calculate precise fitness values for both strains, we measured population frequencies before and after incubation via flow cytometry. We found that small populations of each pure strategist were indeed able to invade majority populations of the other (Figure 2B). Our experimental yeast populations therefore display mutual invasibility between the two pure strategists. Furthermore, there was a unique stable equilibrium frequency of GAL-ON cells that resulted in the same fitness for both pure strategies. Importantly, we find that the frequency of GAL-ON cells that is evolutionary stable is not the frequency that maximizes population growth. Populations with much higher fractions of GAL-ON cells than the equilibrium population grow to saturating density more quickly than the evolutionarily stable population (Figure 2C).

**Figure. 2.**
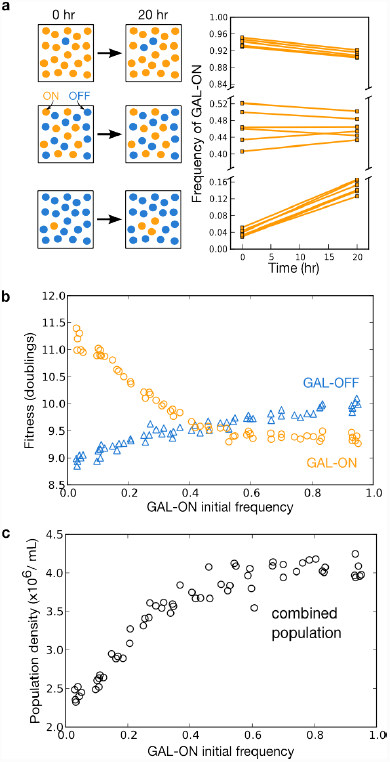
Characterization of the game played between GAL-OFF and GAL-ON pure strategists. **a,** Pure strategists are mutually invasible. Population frequency of the *GAL3*-induced GAL-ON pure strategist (orange circles) relative to the *GAL80*-induced GAL-OFF pure strategist (blue triangles) is plotted at the beginning and end of a 20-hour competition. Six independent cultures of each pure strategist were mixed at high (top panel), intermediate (middle panel), and low (bottom panel) initial frequency of the GAL-ON strain. Each pure strategist invades the other when rare. **b,** Game payoffs (in number of doublings) for both pure strategists are plotted for 60 initial starting frequencies of the GAL-ON strain. The crossing point corresponds to the Nash equilibrium of the pure strategists for the experimental foraging game. **c,** The evolutionarily stable equilibrium is not necessarily growth optimal. Population densities of the mixed populations are shown at 16 hours, before all cultures have reached saturation. Mixed cultures with high initial frequency of the GAL-ON strain grew faster than cultures near the evolutionarily stable mix.

A more in-depth investigation of the dynamics between the pure strategists indicates that the negative frequency dependence is related to the depletion of glucose in the media. Both pure strategists adopt a diauxic growth model; they consume primarily glucose until it is depleted (Figure S2E). Indeed, recent evidence argues that the advantage provided to GAL-ON cells may be primarily due to their ability to consume galactose quickly when the glucose is exhausted ^34^. The GAL-ON pure strategist suffers a fitness disadvantage while glucose is still relatively abundant, but outcompetes the GAL-OFF pure strategist when the glucose becomes low and galactose remains. The GAL-OFF and GAL-ON strategies can therefore be thought of as “specialists” in glucose and galactose, respectively. The hawk-dove game arises because the galactose “payoff” goes to the GAL-ON cells, but the more cells that activate their GAL networks in a population, the slower the glucose gets depleted, and the higher the resulting payoff to glucose specialists (Figure S2).

Game theory predicts that varying the payoff structure of a hawk-dove game correspondingly alters the Nash equilibrium fractions (Box 1). In the context of the simple foraging game, this simply means that if food source A increases, then the stable equilibrium should shift towards a larger fraction of the population specializing in food source A. To test for this phenomenon in the GAL network, we replicated the initial competition of our two pure strategists in eight different concentrations of glucose and galactose. More galactose yields a higher equilibrium fraction of GAL-ON cells, while more glucose yields a lower equilibrium fraction of GAL-ON cells (Figure 3). The stable equilibrium between our pure strategists therefore shifts as predicted by a negative frequency dependent game. However, while this well-behaved shifting of the equilibrium is robust within a range of relatively low sugars, the pattern breaks down in environments of high total sugar concentrations (>0.1%), where carbon may not be limiting in the same way.

**Figure 3.**
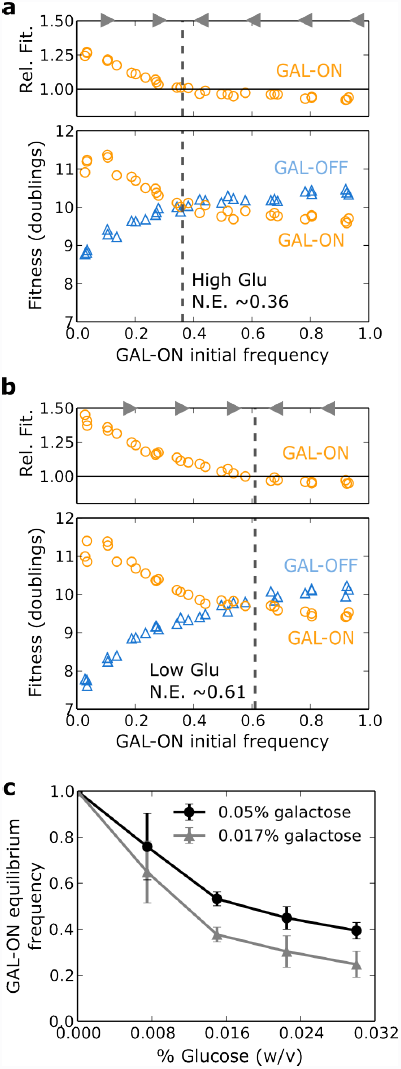
Altering sugar concentration adjusts game payoffs and equilibrium fractions accordingly. **a-b,** Relative fitness of the GAL-ON pure strategist, and absolute fitness (in number of doublings) of both pure strategists is shown for 30 different populations at varying initial frequency of GAL-ON. Data is shown for .05% galactose and two conditions: high glucose (.03%, **a**), and low glucose (.017%, **b**). The payoff for the GAL-ON pure specialists remains roughly the same between the two conditions, while the GAL-OFF pure strategists receive a higher payoff in higher glucose. Lower glucose results in a higher equilibrium frequency of GAL-ON cells, as expected in a hawk-dove like game. **c,** Equilibrium GAL-ON pure strategist frequencies as a function of increasing glucose concentrations. Data is shown for high (0.05%, circles) and low (0.017%, triangles) galactose. All Nash equilibria were calculated by polynomial spline fitting of relative fitness curves (error bars are 95% confidence intervals; n = 3.)

Because the Nash equilibrium mix of pure strategies is a function of sugar concentrations, we next tested whether the wild type mixed cells naturally alter the frequency of mixing based on the concentrations of glucose and galactose. Just as the Nash equilibrium of the pure strategists shifts with varying sugar concentrations, we observed that the mixing frequency of the wild type yeast also shifts: in a higher concentration of galactose, yeast adopt a higher GAL-ON frequency. This type of responsiveness is one of the hallmark predictions of the mixed strategy model. Yeast is able to sense even small differences in the ratio of glucose to galactose and adopts a pure OFF, pure ON, or appropriate mixed strategy accordingly (Figure 4 and S1; also M. Springer personal communication).

**Figure 4.**
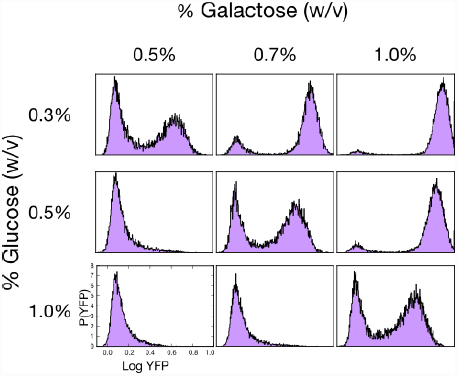
Mixed strategist (wild type GAL network) senses payoff ratio and alters strategy accordingly. The mixing frequency of the wild type mixed strategist is highly responsive to sugar concentrations. GAL network activation level is shown for nine different mixtures of glucose and galactose. From a bimodal expression state, more galactose in the media results in a higher frequency of cells with GAL activation, while more glucose in the media has the opposite effect. This trend is essential to the implementation of an evolutionary stable mixed strategy.

Another prediction of the hawk-dove game is that a strain adopting a mixed ESS cannot be invaded by either pure strategist. However, as the population frequency of the mixed strategist approaches one, it becomes only neutrally uninvasible. In other words, in the limit of a population consisting entirely of mixed strategists, any single invading cell adopting any strategy (pure or mixed) will receive a payoff equal to the mixed strategist. By competing the pure strategist strains (GAL-ON/OFF) with a strain containing the wild type GAL network (mixed strategist), we determined that the mixed strategist is indeed uninvasible by either pure strategist. Additionally, a competition between pure GAL-OFF and the mixed strategist displays the neutral uninvasibility predicted from the game theoretic model (Figure 5A). The wild type mixed strategy can spread in a population of GAL-OFF cells, but as the wild type strategy increases in frequency, its advantage disappears. Moreover, the wild type mixed strategist cells are uninvasible by the GAL-ON pure strategist at all frequencies (Figure 5B), though the interaction does not display strong frequency dependence. This lack of strong frequency dependence between this pair suggests that the dynamics of yeast in mixed sugar environments have some subtle deviations from a simple foraging game.

**Figure 5.**
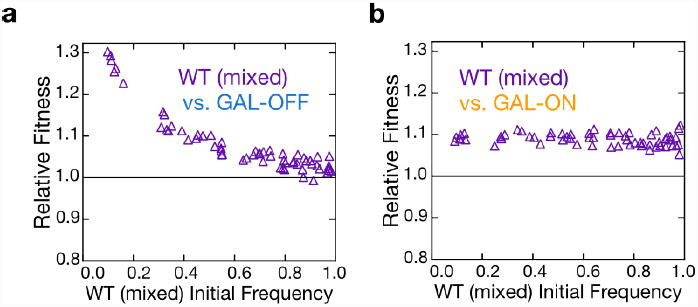
Wild type mixed strategist invades both pure strategists and is uninvasible by either. Relative fitness of the wild type mixed strategist over the GAL-OFF pure strategist (**a**) and GAL-ON pure strategist (**b**) is shown. Low frequencies of the mixed strategist invade strongly in populations dominated by either pure strategist. As expected of an evolutionarily stable mixed strategy, the relative fitness of the mixed strategist to the GAL-OFF pure strategist approaches one in populations dominated by the mixed strategist. However, the mixed strategist does not display frequency dependence against the GAL-ON pure strategist.

### Discussion

When observing phenotypic heterogeneity in microbial populations, it is important to consider the underlying evolutionary reasons for heterogeneity and distinguish between the different explanations where possible. While it is very difficult (if not impossible) to prove claims about historical reasons for the evolution of stochasticity in specific systems like the yeast GAL network, different evolutionary drivers of heterogeneity do make unique and experimentally verifiable predictions about the fitness dynamics between the associated phenotypes (see Table 1 and deJong and Kupers’ review^18^). In this work we have demonstrated a simple way of probing whether observed phenotypic heterogeneity might be implementing an evolutionarily stable mixed strategy. By isolating the pure strategies and probing them for mutual invasibility, we have determined that a hawk-dove style foraging game is being played between the GAL-ON and GAL-OFF strategy. This frequency-dependent mutual invasibility distinguishes a mixed ESS from bet-hedging (which does not rely on interactions within the population) and altruistic division of labor (in which the altruistic phenotype is always less fit). We have also verified the theoretical prediction that the evolutionarily stable mixed strategy is not necessarily optimal for growth, and confirmed that a strain implementing a mixed strategy invades populations of both pure strategists, and is uninvasible by either.

In the mixed sugar conditions we have shown, the wild type mixing frequency is roughly the same as the Nash equilibrium between the mutant pure strategists (compare figure 1d with figure 2b). However, we do not expect the quantitative agreement to be general, since budding yeast did not evolve its mixing frequency in laboratory cultures of mixed glucose and galactose. There are also slight differences between the mutants and the wild-type phenotypes. For example, the GAL-repressed subpopulation of the mixed strategist adopts a diauxic growth phenotype: it activates its GAL network upon glucose depletion (Figure S3), but because of the doxycycline induction of *GAL80*, the GAL-OFF pure strategist does not transition to GAL-ON within the time frame of the competition (Figure S3). Consequently, in a mixed sugar scenario, the GAL-suppressed fraction of the mixed strategist is likely to be more fit than the GAL-OFF pure strategist. Additionally, the GAL-ON pure strategist’s induction activates the GAL network to a greater degree than the induction in the wild type (Figure 1D), resulting in slightly different costs for expressing the GAL network. Also, while the majority of the wild type yeast’s stochastic decision to be GAL-OFF or GAL-ON is determined early, there is a small amount of stochastic switching between the states (Figure S4), which does not occur in the mutant pure strategists.

Negative frequency dependent interactions are often invoked as reasons for stable coexistence, and evolutionary stable mixed strategies (in the context of hawk-dove games) are central to evolutionary game theory. Yet this broad class of interactions has received almost no attention as an evolutionary reason for phenotypic heterogeneity in clonal populations. To our knowledge this work constitutes the first experimental evidence that phenotypic diversity in an isogenic microbial population is—at least in part—implementing a game theoretic mixed strategy in response to a negative frequency dependent foraging game. It remains to be seen to what degree such games are responsible for the widespread phenotypic heterogeneity in isogenic populations.

### Materials and Methods

*Strains:* The three strains of *Saccharomyces cerevisiae* (wild type mixed strategist, GAL-OFF specialist and GAL-ON specialist), are modified from those used in Acar et al. (2005), which were derived from the diploid W303 strain of *S. cerevisiae.* All strains have a ADE2-P_GAL1_-YFP reporter construct inserted at one *ade2* site for monitoring activation of the GAL network. Since one *ura3* locus was already occupied by inducible forms of *GAL80* or *GAL3*, yeast was first sporulated to isolate the remaining *ura3* locus. Identity of the haploids was confirmed by replica plating. Haploids containing *ura3* were then transformed with the yeast integrating vector pRS306 containing URA3 and either RFP(tdTomato) or CFP cloned downstream of a TEF1 promoter. Constitutive fluorescence was confirmed by microscopy and flow cytometry. Fluorescent cells were then mated with the appropriate haploid to produce the desired strain. All strains were maintained on synthetic media his-and ura-agar dropout plates supplemented with 2% glucose.

The Gal80-inducible (GAL-OFF pure strategist) strain has a double *GAL80* deletion. *P_TETO2_-GAL80* is inserted at one *ura3* locus, while *P_MYO2_-rtTA* is inserted at an *ade2* locus. The GAL3-inducible (GAL-ON pure strategist) strain has a double *GAL3* deletion with P_TETO2_-GAL3 inserted at a *ura3* locus and PMYO2-rtTA inserted at an *ade2* locus. Complete genotypes for the strains are found in the Supplementary Information.

*Competitions*: To initiate doxycycline induction in pure strategists, strains were initially mixed at desired initial frequencies from plated colonies, then incubated in 1.0 µg/mL doxycycline and 0.01% (w/v) glucose for 24 hours from a starting density of ~3x10^4^ cells/mL to a saturating density of ~6x10^6^ cells/mL, then diluted to ~3x10^4^ cells/mL in synthetic media supplemented with glucose and galactose as indicated. Fractions were measured before and after incubation using a Miltenyi MACSquant flow cytometer (20,000+ cells per well), and population density was measured as absorbance at 600 nm in a microplate spectrophotometer (conversions assume 3×10^7^ cells/mL at A_600_ = 1.0).

## Acknowledgements

The authors thank A. Sanchez, A. Velenich, L. Gibson, M. Springer, M. Laub and M. Vander Heiden for helpful discussions, K. Axelrod for experimental help, and the members of the Gore Laboratory for helpful comments on the manuscript. M. Acar provided the strains from which the current strains were modified. This work was funded by the Allen Distinguished Investigator Program, NIH New Innovator Award, and NSF CAREER Award. JG also thanks the Pew Scholars in the Biomedical Sciences and the Sloan Fellows Program.

## Author contributions

D.H. and J.G. designed the study and performed analysis. D.H. performed the experiments. D.H. and J.G. wrote the manuscript.

